# Mapping wild bee diversity and flower use for an effective conservation of the Doñana biodiversity hotspot

**DOI:** 10.1101/2025.10.29.685282

**Authors:** Francisco P. Molina, Nerea Montes-Perez, Luis Villagarcía, Ignasi Bartomeus

## Abstract

Despite the high bee species richness found in the Iberian Peninsula, the lack of distributional data often hampers effective conservation. This gap is particularly critical within protected areas such as the Doñana Natural Area, where biodiversity is expected to be increasingly vulnerable to environmental pressures. Here, we present the first checklist of the wild bees of Doñana. A total of 385 species belonging to 47 genera were recorded, including recently described species and a newly recorded species for continental Europe, *Andrena purpurascens*. The exceptional diversity of the area’s bee fauna accounts for approximately one-third of the known Iberian apifauna. We provide information on habitat and floral use, offering key ecological insights for conservation planning. This knowledge is essential to support habitat management decisions, particularly in the face of intensifying climate change and water scarcity across the protected area. In addition, to evaluate long-term ecological changes, we compared recent field data (2020–2021) with historical surveys conducted at the same site between 1984 and 1985. While a similar number of species were recorded in both periods (49 vs 43 species), only 15 species were shared between the two periods. This large-scale species turnover indicates the dynamic nature of a region undergoing significant environmental shifts. These findings underscore the importance of baseline biodiversity assessments and long-term monitoring for understanding and mitigating pollinator loss in protected ecosystems, which are increasingly shaped by climate instability and anthropogenic pressures.

## Introduction

The Doñana Natural Area, located in southwestern Spain and spanning the provinces of Huelva, Cádiz, and Seville, encompasses both a national and a natural park. It is one of the most emblematic protected areas in Europe, notable for its exceptional biodiversity and ecological significance (Berta et al,. 2011). Recognized as a UNESCO Biosphere Reserve since 1981 and a World Heritage Site since 1994 (UNESCO, 2024), Doñana supports a wide variety of habitats shaped by its sandy and clay substrates, Mediterranean climate, and highly variable rainfall patterns.

While Doñana is well known for its vertebrate fauna, including iconic species such as the Iberian lynx (*Lynx pardinus*) and Spanish imperial eagle (*Aquila adalberti*), arthropods, and pollinators in particular, have received comparatively little attention. Despite the ecological relevance of bees (Hymenoptera: Apoidea: Anthophila) as key pollinators in Mediterranean systems (Ollerton et al., 2017; Winfree, 2010), no comprehensive inventory of bee species has been published for this area to date. Existing records are scattered, limited in taxonomic and spatial coverage, and often derived from non-systematic surveys (Herrera, 1988; Talavera et al., 1988; Magrach et al., 2021). This lack of baseline information prevents any assessment of long-term changes and limits the development of conservation strategies targeted at this critical functional group (Noriega et al., 2018). Moreover, ecological information such as bee–flower interactions is virtually absent, even though these interactions are essential for understanding plant–pollinator network structure, floral preferences, and pollinator specialization levels. Documenting these associations is not only critical from a theoretical ecological standpoint (Bascompte et al., 2006) but also for identifying key plant resources that support pollinator diversity (Blaauw and Isaacs, 2014).

Understanding bee floral use and preferences is essential for conservation because it not only informs bee species food requirements but also structures the web of interactions that shapes community assembly and supports population persistence (Valdovinos et al., 2016; Domínguez-Garcia et al., 2024). Moreover, disturbances like climate change are rewiring plant–pollinator networks, making informed management vital to prevent cascading consequences (Burkle et al., 2013). In fact, bees are known for being generalist foragers (Waser et al, 1996). Although some loose trait matching is expected (e.g., long-tongued bees on long-corolla flowers; Stang, 2009), floral resource abundance and diversity modulate visitation rates (Laha et al., 2020). Behavioural flexibility is common and generalist species can switch flower species depending on floral resource availability (Papaj and Russel 2024; Fründ et al., 2010). Finally, competitive contexts further modulate realised floral use; for example, competition among pollinators, which can be intensified by managed honeybees in some systems, can reshuffle species interactions (Magrach et al., 2017; Sponsler et al., 2024; Requier et al., 2024). Yet, despite the importance of understanding floral use by pollinators, it remains unquantified in most Mediterranean systems, including the Doñana Protected Area.

In this study, we compile and integrate over a decade of systematic fieldwork and four decades of opportunistic observations, to generate the first species-level checklist of wild bees for the Doñana Protected Area. Our sampling encompasses a broad range of habitats, including Mediterranean scrublands, marshlands, and dune-beach systems. It captures both taxonomic and ecological data, linking wild bee species to the floral resources they visit. Beyond compiling species occurrences, we also analyse (i) floral usage patterns, (ii) the relative importance of plant species to bee communities, and (iii) temporal changes in bee composition by comparing recent data to historical records from the same site. Overall, we expect that shrubs are the most important plants for pollinators, and a shift in bee species dominance over time with a potential loss of species linked to environmental pressures. In doing so, we provide not only a biodiversity baseline for a poorly known pollinator community, but also a window into how the community and its ecological interactions may have changed over time. The approach taken here combines species and interaction data across decades, contributing new insights into Mediterranean pollination networks and informing conservation strategies in one of Europe’s most iconic protected areas.

## Materials and Methods

This study was conducted within the Doñana Protected Area (southwestern Spain), which spans 128,386 hectares and encompasses a wide variety of habitats. Among the most ecologically relevant are the Mediterranean forest and scrublands (*cotos*), the ecotonal zone known as *la vera*, and the coastal dune-beach system. The distinct plant communities associated with each of these environments provide diverse floral resources that support a rich wild bee fauna.

### Habitat Characterization

The Doñana Protected Area is a unique ecosystem shaped by the convergence of diverse habitat types, resulting in high levels of biodiversity. For this study, we grouped the sampling sites into three major habitat categories to facilitate ecological interpretation:

- **Marshland**: This is the most extensive habitat in Doñana, characterized by pronounced seasonality and hydrological fluctuations linked to rainfall. The marshes support vegetation adapted to wet or saline conditions, including plant species such as *Eryngium maritimum*.
- **Dunes and beaches**: This coastal system comprises active dunes, sandy soils, and beach environments, forming a transitional zone between marine and terrestrial ecosystems. Its flora includes sandy-soil specialists such as *Lotus creticus*, *Malcolmia maritima*, and *Helichrysum picardii*, all of which are important nectar and pollen sources for many bee species.
- **Mediterranean forest and scrub (*cotos*)**: This habitat features extensive pine (*Pinus* spp.) woodlands interspersed with diverse shrub layers. It supports over 30 flowering species, particularly from the Lamiaceae (e.g., *Lavandula pedunculata*, *Salvia rosmarinus*, *Thymus mastichina*) and Cistaceae (e.g., *Cistus salviifolius*, *Cistus libanotis*, *Halimium halimifolium*) families, along with Ericaceae such as *Erica ciliaris*, *Erica umbellata*, and *Calluna vulgaris*.

### Data collection

In the absence of a historical reference collection for wild bees in Doñana, we compiled early records from entomologists and research projects to reconstruct a baseline of the region’s bee fauna. The earliest available survey was conducted between 1983 and 1984 by F.J. Herrera Maliani (University of Seville), who documented pollinators and their floral resources at a single location—Laguna del Ojillo—within the Doñana Biological Reserve. To enrich this historical dataset, we also incorporated digitized specimens from opportunistic collections by Klaus Warncke, Javier Ortiz, Stuart Roberts, and Carlos M. Herrera, currently housed in the Iberian Bees database (Bartomeus et al., 2022).

Building on this historical foundation, a long-term monitoring program was established between 2015 and 2025, encompassing 17 scrubland sites, including the original site at Laguna del Ojillo. At each site, bee–flower interactions were recorded using two complementary methods. First, standardized 30-minute transect walks were carried out along 100-meter paths and repeated at least seven times annually throughout the flowering season. All flower-visiting bees that could not be confidently identified in the field were captured with hand nets, euthanized via freezing, and stored in labelled vials. Specimens were later pinned and identified in the laboratory using morphological keys and reference collections, following established wild bee sampling protocols (Magrach et al., 2021; Domínguez-García et al., 2024).

In parallel, focal observations were conducted during the same sampling rounds on dominant shrub species, *Cistus salviifolius, C. crispus, C. ladanifer, Lavandula stoechas, Salvia rosmarinus, and Teucrium fruticans*. For each plant species, three individuals per site were monitored in 3-minute observation periods. All insect visitors were collected using the same methods applied during transect walks (Tobajas et al., 2023).

To complement these systematic efforts and address spatial gaps in historical records, targeted surveys were conducted over the past decade in areas that had previously received little or no sampling effort. Particular attention was given to peripheral zones and underrepresented or logistically challenging habitats such as dense scrublands, mobile dunes, and seasonal wetlands. These habitats, although often overlooked, are known to harbor unique and specialized bee assemblages that may be absent from more accessible areas. As such, non-systematic sampling played a critical role in uncovering previously undocumented species and expanding both the taxonomic and ecological breadth of the Doñana bee inventory. For instance, bee species such as *Osmia rutila* and its cleptoparasite *Dioxys ardens* were recorded in beach and dune habitats that had not been previously surveyed within the study area. The value of these exploratory surveys has been highlighted in recent literature as essential for capturing rare or cryptic taxa, improving biodiversity assessments, and informing conservation planning in heterogeneous landscapes (Hending, 2024).

### Taxonomic identification

Most collected specimens were linked to their visited plant species, allowing insights into floral associations.

For flora, specimens were pressed using a botanical press, and species were identified using specific bibliographic references, notably: Valdés Castrillón, Benito; Talavera Lozano, Salvador & Fernández-Galiano Fernández, Emilio (1987), and Bonnier, G., & De Layens, G. (1990). Additionally, several experts contributed to the identification of certain specimens (see acknowledgments).

Bee species were primarily identified by the first author of this study. All specimens were identified to species level and, where possible, to subspecies level. An extensive range of bibliographic resources was consulted, including Amiet et al. (2001, 2004, 2007, 2010, 2014), Aubert (2020), Bogusch & Straka (2012), Blüthgen (1924), Dardon et al. (2014), Kasparek (2015, 2022), Michez et al. (2004), Ornosa & Ortiz-Sánchez (2004), Ornosa et al. (2009), Ortiz-Sánchez, F. J., & Jiménez-Rodríguez, A. J. (1991), Ortiz-Sánchez et al. (2003, 2004, 2009, 2012), Pauly (2016), Schmid-Egger & Scheuchl (1997), Smit (2018), Terzo & Ortiz-Sánchez (2004), Terzo & Rasmont (2007), Verges (1967), and Wood TJ (2023). In addition, several experts were consulted for the identification or confirmation of specimens that presented taxonomic difficulties (see Contributors and Acronyms section). Newly described species, such as *Andrena baldocki* (Wood, 2024) or *A. ramosa* (Wood et al., 2022), are also included in this inventory following their primary descriptive paper (Wood, 2024). For already collected material, we used the original taxonomical identification as indicated in Supplementary S1 (see below Contributors and acronyms section).

### Data cleaning and checklist creation

We selected records for which we had full information on the bee species name, coordinates or location, and date. After performing these filters, we end up with a final dataset of 28.666 total records. Taxonomic information was checked and corrected following the taxonomic classification by Ortiz-Sánchez (2020a). Additionally, data with plant visitation information was also revised to ensure plant names follow the Catalogue of Life classification. Data were summarised by bee species, site, date, and visited plant species when possible, obtaining the total number of females, males, or workers per observation. Records with no sex assigned were given the category of “unknown” to avoid losing information, as 7.2% of records did not have sex information.

With the final dataset, we created a checklist of species for the Doñana Protected Area. The complete taxonomic information for all specimens was transferred to a database in a standardized format, from which the species inventory was generated. The tribes and genera within each subfamily, as well as subgenera and species, are organized in alphabetical order. In the species list, the locality or site where the specimen was collected is mentioned first, followed by the collection date. For each locality and date, the number of collected specimens and their sex (if available) are specified. Then, in brackets, the name of the collector (leg.) and the person who identified the specimens (det.) are indicated.

### Contributors and acronyms

The acronyms used for the authors of this work are as follows: Francisco de Paula Molina Fuentes [FPM], Nerea Montes Pérez [NMP], Alejandro Núñez Carvajal [ANC], Denis Michez [DM], Félix Torres González [FTG], Guillaume Ghisbain [GG], Luis Oscar Aguado [LOA], Simone Flaminio [SF], and Thomas J. Wood [TW]. Non-systematic observations collected in the Iberian Bees database were checked by several taxonomists, whose full list is provided at the end of Supplementary S1.

### Statistical analyses

First, we assess species completeness by using accumulation curves based on rarefaction analysis using the package iNEXT (Hsieh et al., 2016). Second, we focused the analyses on two aspects: the floral usage of wild bee species and the historical trends of the wild bee community inside the National Park. From the 28666 total records, 97% of records contained information about plant species, lacking plant data for only 14 bee species. Results for those 14 species are shown as unknown in all floral analyses.

For the floral analysis, we obtained the bee plant usage and calculated the plant usage and importance. Bee plant usage was described per species, taking into account species’ known behaviour. Thus, we calculated floral usage in five different levels: generalist (usage of multiple families), family-usage (visits plants only from one family), genus-usage (visits plants from only one genus) and species-usage (only visits one plant species), and unknown (no data available). We categorized each bee species into one of those five usage groups, looking at their visitation patterns. We assigned a category when 95% of visits fall into one of these categories. In Supplementary S2 Table S1, it can be found the set of plant genera visited by each bee species. This categorization is based solely on the data collected in the Doñana Protected Area and reflects the observed behaviour and floral resource use of the listed bee species within this context. It is not meant to characterize the full ecological breadth, degree of specialization, or floral preferences of each species across its entire range.

Plant importance *sensu* Bascompte et al. (2006) was calculated taking into account the number of bee species that visited each plant and the floral dependency of each bee to that plant. Bee dependency is calculated as the number of visits to a plant species divided by the total number of visits of the bee. This metric reflects high values for plants supporting many species or supporting species highly dependent on those plants.

Lastly, we compared the field sampling of the Ojillo lagoon area, located within the core protected zone of Doñana (RBD-CSIC), during 2020 and 2021, with the data collected by Professor Javier Herrera (Faculty of Biology, University of Seville) at the same site during 1984 and 1985. All specimens in both periods were identified to the species level, allowing for direct comparisons of the species identities across decades. Unfortunately, no flower usage data is available for the older time period. We assessed community changes between the data from the sampling in 1983 -1984 and the sampling in 2020 -2021. We used the R package *betapart* to calculate composition changes between past and present and understand which percentage of change is due to turnover (species replacement) or nestedness (species gain/loss). Composition changes were obtained using Sørensen dissimilarity (Baselga et al., 2010).

All data cleaning, statistical analyses, and figures were done using software R (v 4.5.1), and figures were done using package *ggplot* (Wickham, 2016), *ggalluvial* (Brunson, 2020), *eulerr* (Larsson & Gustafsson, 2018), and *circlize* (Gu et al., 2014).

## Results

### Composition of Bee Fauna in the Doñana Natural Area

The Doñana Natural Area represents one of the most ecologically significant regions in southern Europe, and its rich assemblage of pollinating Hymenoptera—particularly bees (Apoidea: Anthophila)—is a clear indicator of its biological diversity. To date, a total of 385 bee species have been identified within the area, spanning 47 genera and belonging to all six families recognized in the Iberian fauna: Andrenidae, Apidae, Colletidae, Halictidae, Megachilidae, and Melittidae (Table 1). The richest family is the Andrenidae family, reaching 100 bee species, followed by the Apidae family with 98. Among all families, approximately 14% of species are cleptoparasitic species. The proportion of eusocial taxa cannot be accurately estimated, since the nesting and social behavior of the majority of species remain insufficiently documented.

**Table 1:**
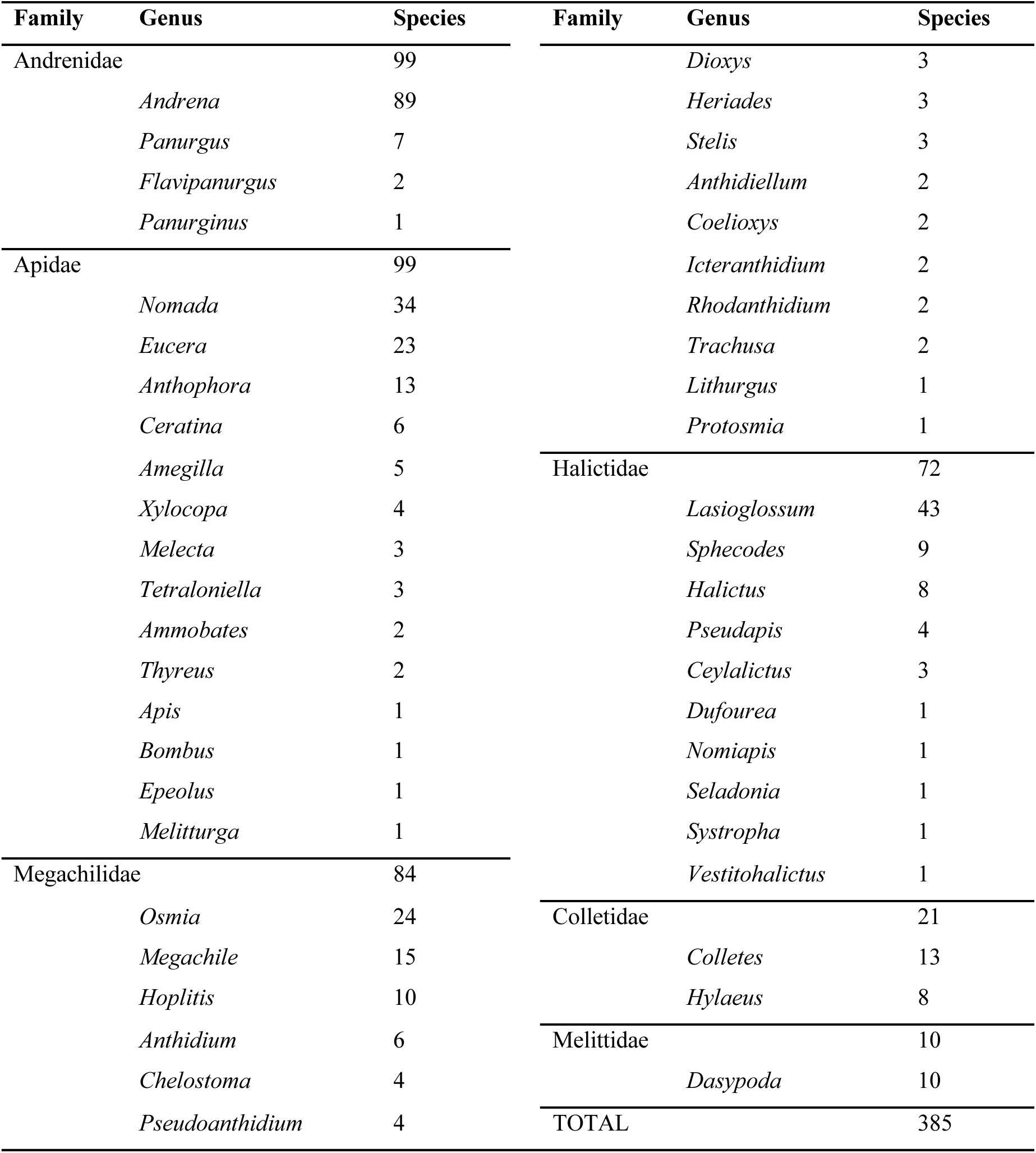
Bee species diversity summary per family and genus. Families and genera are ordered by species richness, showing the most abundant first.

A notable outcome of our inventory is the detection of *Andrena purpurascens* (Pérez, 1841) in the Doñana Protected Area (SW Spain), which represents the first confirmed record of this species for the Iberian Peninsula and continental Euro-e. This bee, assigned to the subgenus *Distandrena*, was previously known only from North Africa, with published records from Morocco, Tunisia, and Algeria. Its occurrence in Doñana marks a significant range extension and may reflect a biogeographical link between southern Spain and North Africa, but we cannot know if the species’ presence represents a recent colonization. Photographs of the specimen can be found in Supplementary S2 Figure S1. This finding underlines the value of comprehensive faunistic surveys in Mediterranean biodiversity hotspots such as Doñana. The complete annotated checklist can be found in Supplementary S1.

These results reflect not only an outstanding level of species richness but also the ecological uniqueness of the Doñana natural area. In comparative terms, Doñana harbors approximately one-third of the approximately 1170 bee species known from the Iberian Peninsula (Ortiz-Sánchez, 2020; Reverté et al., 2023), and nearly two-thirds of those estimated for the region of Andalusia. This highlights the site’s critical role as a refuge for the conservation of wild bees and other pollinators. As expected, the managed *Apis mellifera* is the most abundant species, followed by *Xylocopa cantabrita, Bombus terrestris,* and *Lasioglossum immunitum* (Figure 2(a)). Interestingly, only 9% of bee species have more than 100 individual records, while 76% have 20 or fewer observations, with 25% represented by a single recorded individual. These findings highlight the significant imbalance in the dataset, with most bee species being rare or known from limited records.

**Figure 1:**
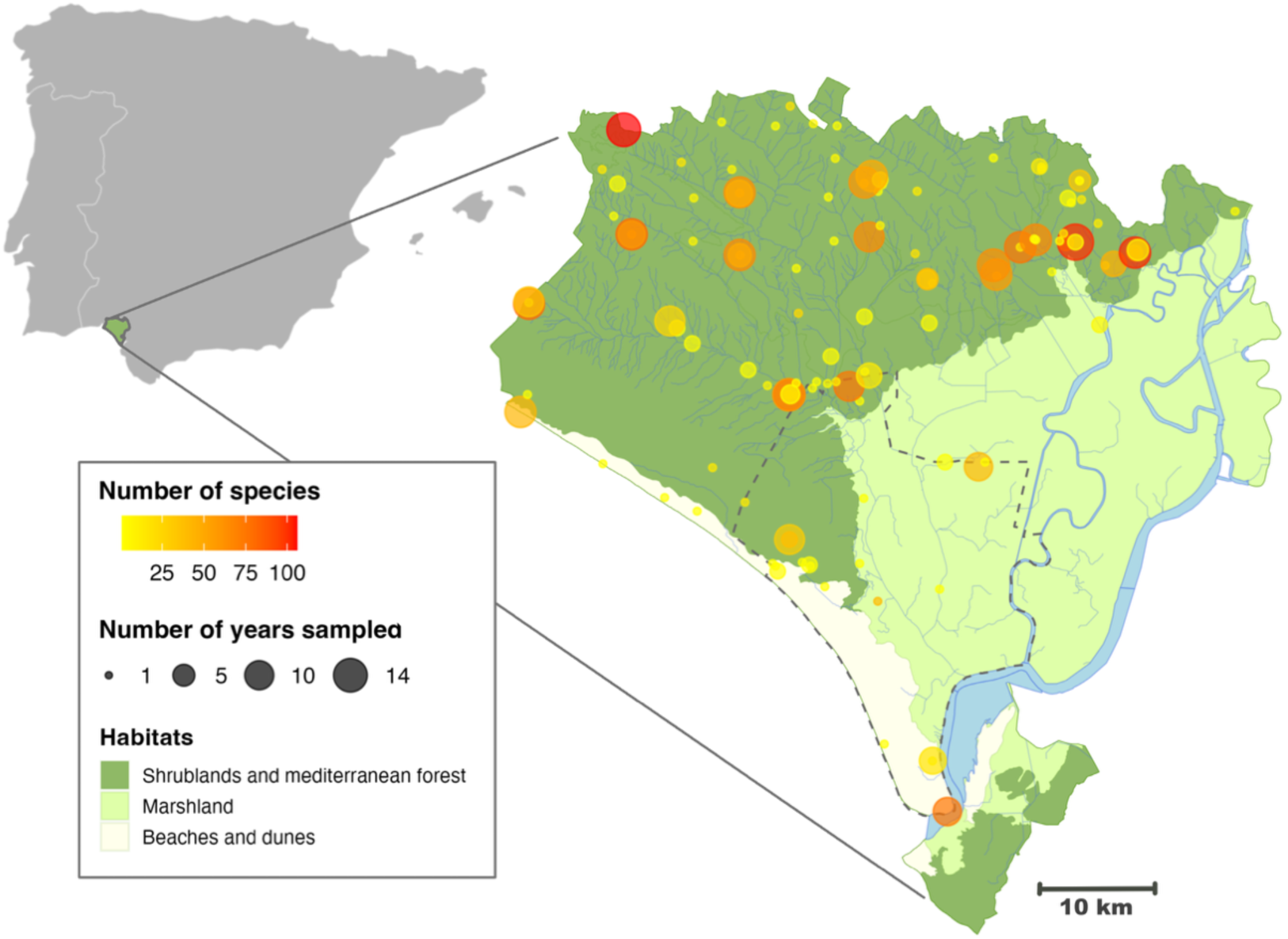
Map of the sampled areas inside the Doñana Protected Area. Dots represent sampling sites, with dot size representing the number of years it was sampled and with colour the number of species found per site (yellow to red showing low to high number of species). The area is divided into three main habitats: shrublands and Mediterranean forest (the most abundant), followed by marshlands, and finally beaches and dunes. All data collected from the study area have been included in the map.

**Figure 2:**
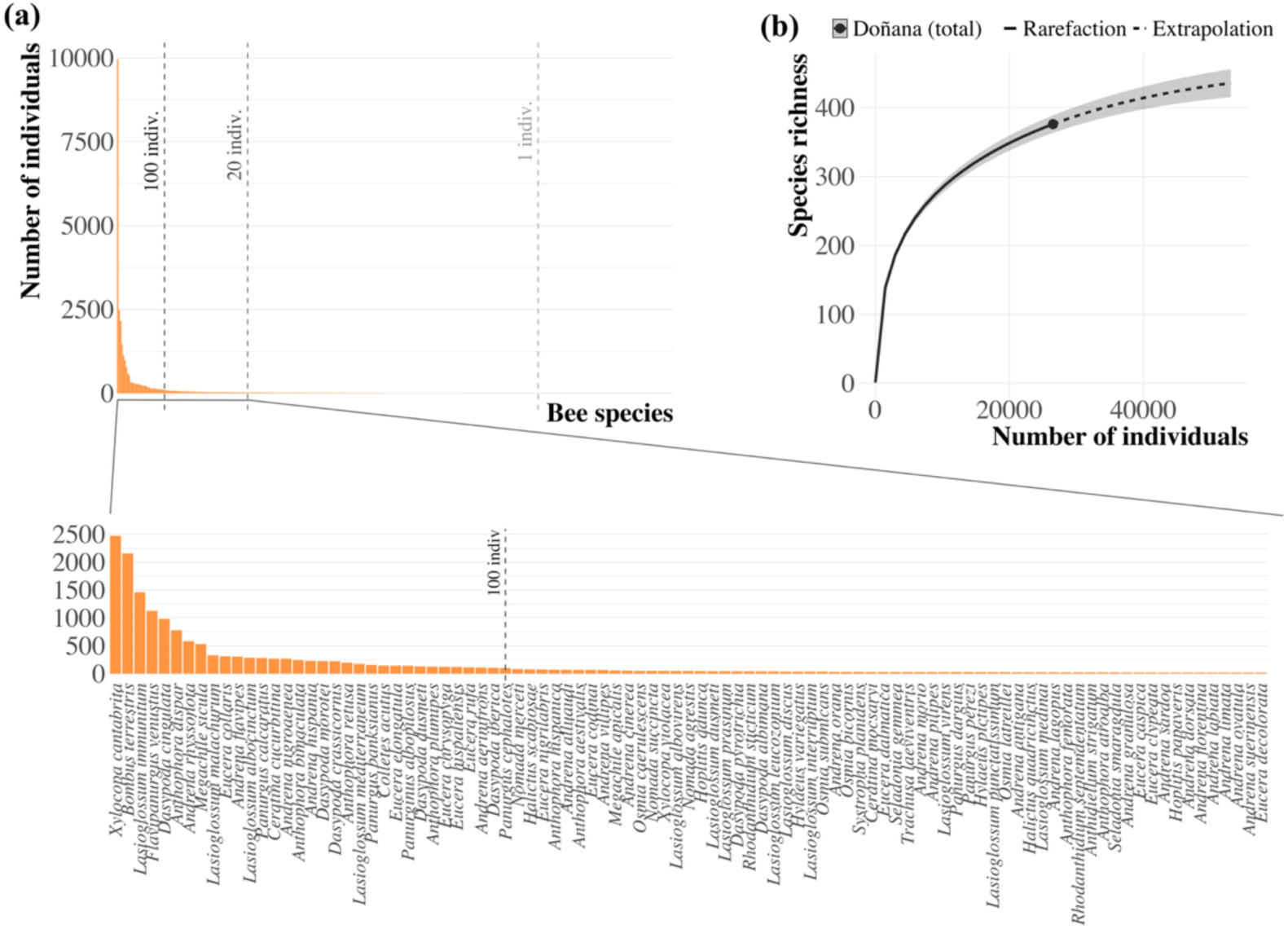
(a) Bee species abundance for all bee species collected in this inventory. Marked with dashed lines from left to right are the limits of species with more than 100 individuals, species with more than 20 individuals, and species with only 1 individual recorded. The most abundant species is *Apis mellifera*. At the bottom, a close-up of all species with more than 20 individuals, starting with the second most abundant. (b) represents the rarefaction curve for the Doñana Protected Area based on sampling size.

Accordingly, species accumulation curves based on rarefaction analyses (see Figure 2(b)) indicate that, although we have sampled a large fraction of the species (Coverage = 99%), sampling has yet to reach an asymptote, suggesting that additional species likely remain undetected and that the actual richness may still be underestimated.

### Bees’ floral usage

In total, 371 bee species have flower visitation data available, encompassing interactions with 147 different plant species. Thus, we have been able to obtain the pollination interaction network for the Doñana Protected Area for 97% of the total bee species recorded. This network of plant-pollinator interactions shows a core of abundant species interacting among each other, and many rare species, mostly interacting with the core (Figure 2). The list of plant species can be found in Supplementary S2 Table S2.

From those 385 bee species, 236 (61%) visit multiple plant taxa (i.e., they interact with plants from different families) and 120 (31%) use a single plant species. Note that, out of those bee species that visited a single plant species, 93 (77%) were recorded only once, and hence, their diet cannot be reliably assessed. If we look at the distribution of bees depending on the breadth of their visitation pattern across bee families, Melittidae is the only family that does not have bees that are single-plant species visitors (Figure 3, Table 1).

**Figure 3:**
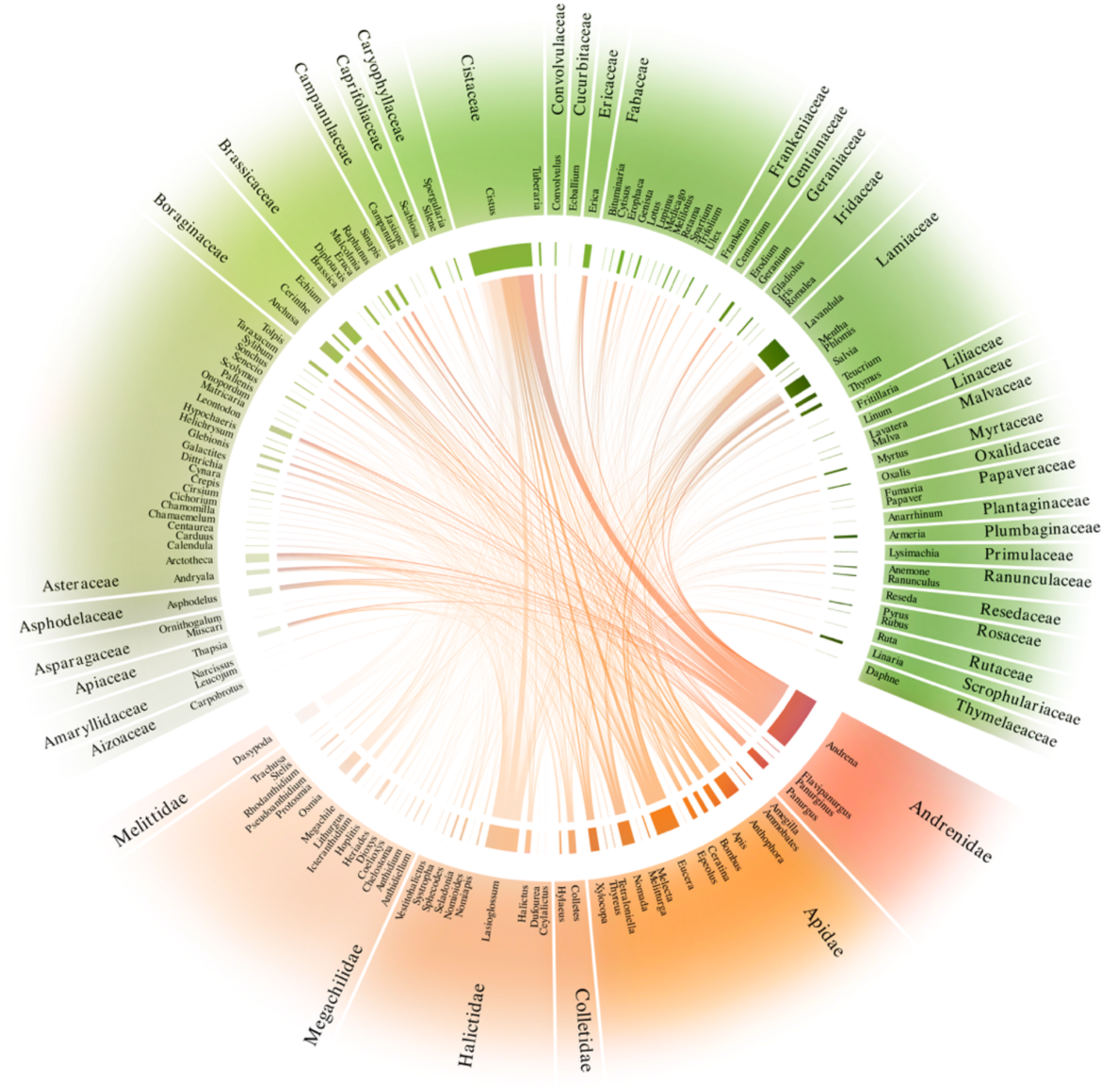
Summary of the interaction network by genus and family of all wild bee species. Based on 28093 records with plant data out of the 28666 total records collected in this dataset. Pollinator families and genera are represented with a range of orange colours, and plants with a range of green colours. The most visited plant genera are *Cistus* and *Lavandula*, which comprise 7 and 3 species, respectively. In the case of pollinators, the genus with the greatest number of unique interactions is *Andrena*, which is the most diverse wild bee genus with 89 species.

Looking at the visited plants, we found that species from the families Cistaceae and Lamiaceae are the most important ones in terms of wild bee species that visit them (Figure 4 (a)). Among these species, it is important to highlight *Cistus crispus*, *Salvia rosmarinus,* and *Cistus salviifolius,* which provide floral resources for over 100 bee species each.

**Figure 4:**
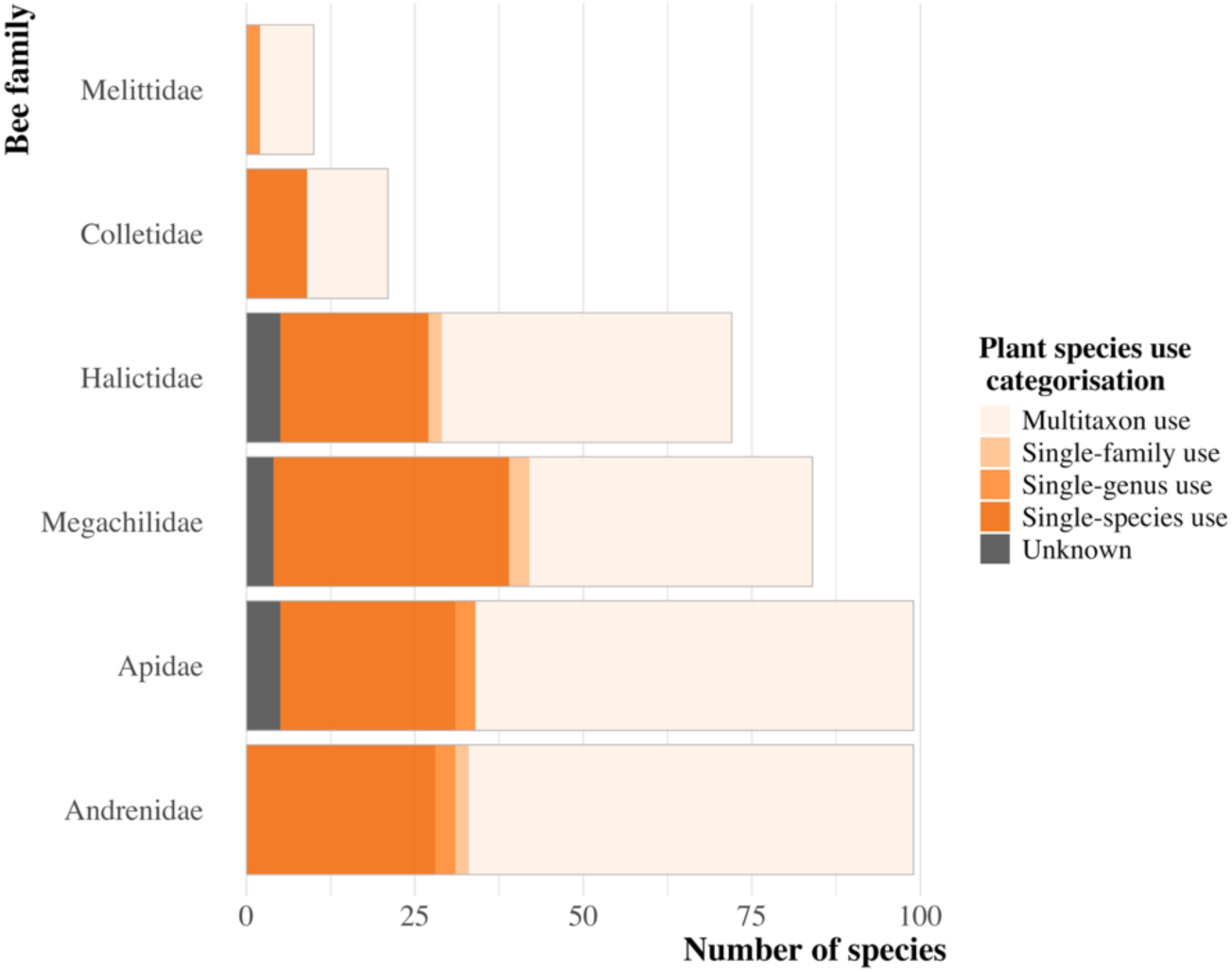
bee species categorization depending on plant species use per family. In an orange gradient, from multitaxon use (the species visited multiple plant families), single-family use (only visit one plant family), singles-genus use (only visit one plant genus) and single-species use (only visit one plant species). In grey, species for which we do not have plant visitation data.

When we assessed plant importance using floral dependency of pollinators, we found similar results, where the most important plants from Figure 4 (a) are also those with a higher number of bee visitors (Figure 4 (b)). There is a positive correlation of 0.91 ± 0.03 between the total floral dependency (i.e., importance) of a plant and the number of bees visiting a single plant this plant sustains. Thus, protecting the most important plants ensures keeping floral resources also for many bees with high dependence values. It is important to stress that there are some examples of plants visited only once, such as *Papaver rhoeas* or *Ornithogalum baeticum,* whose unique interaction is with a single-species visitor. The summary list of bee preferences can be found in Supplementary S2 Table S1.

### Historical trends

A total of 49 wild bee species were recorded in the recent surveys, compared to 43 species documented in the historical dataset (Figure 5 (a)). Despite this relatively similar species richness, only 15 species were shared between the two periods (Figure 5 (a) and (b)).

**Figure 5:**
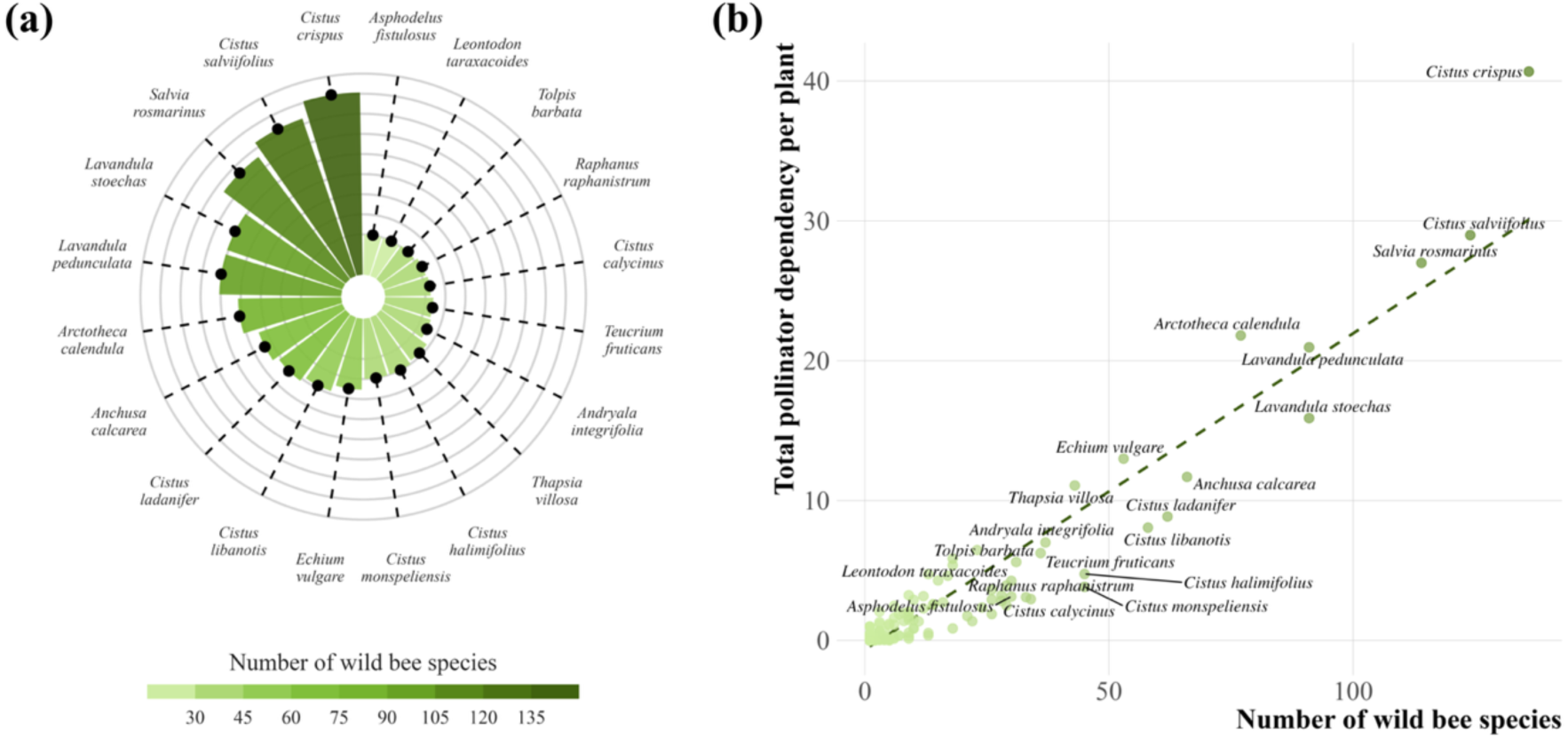
(a) The 20 plant species most visited within the Doñana Protected Area, where the darker the colour, the more bee species visited it. (b) floral importance for all plant species. Plants above the dashed line have a larger importance than expected, given the number of floral visitors, because they sustain dependent pollinators. Colours match the species visitation gradient of plot (a), and labels are assigned to those plant species visited by more than 30 bee species.

**Figure 6:**
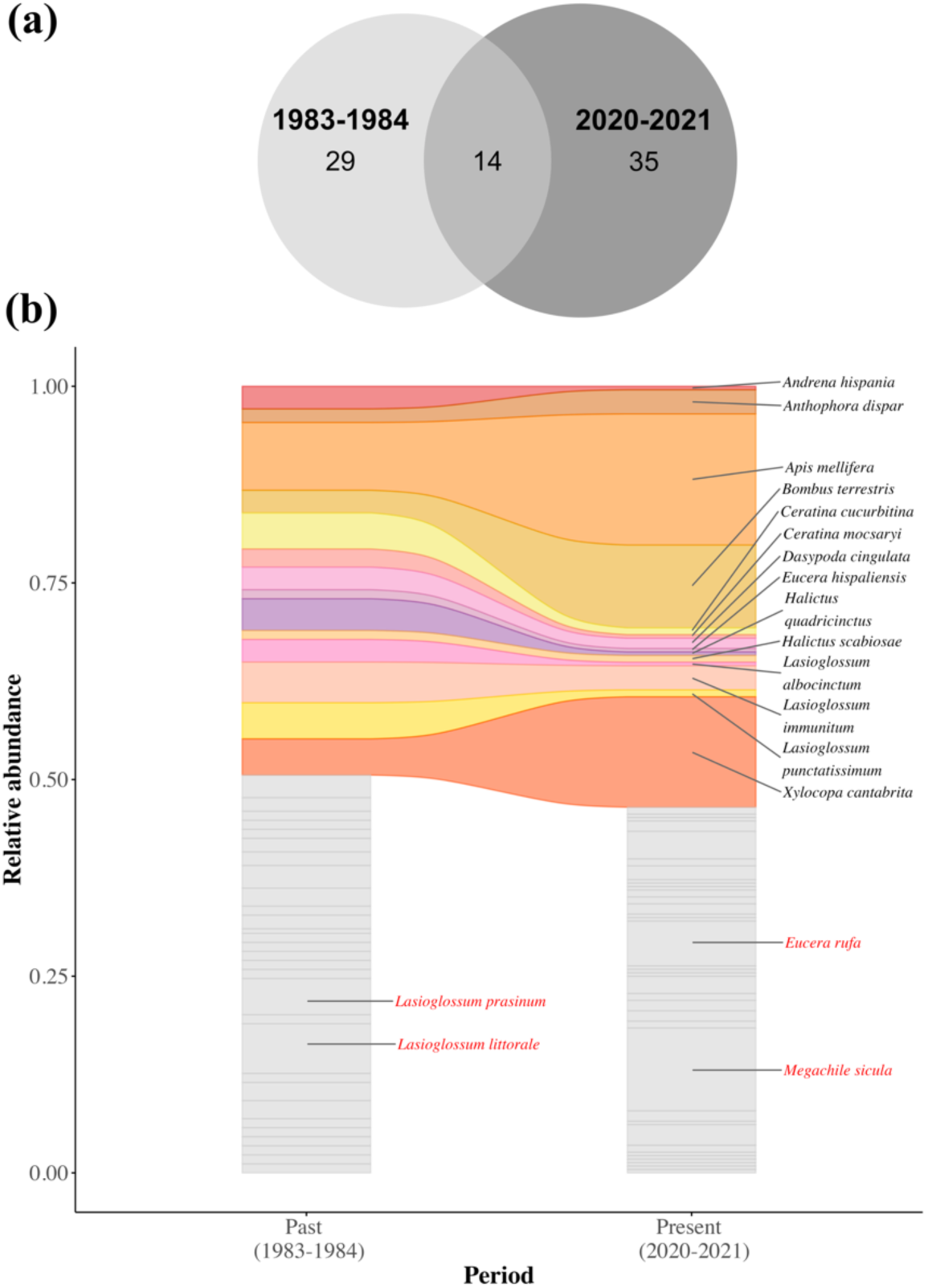
(a) Venn diagram showing the number of species per period and the total shared, and (b) alluvial plot showing the relative abundance of species in both periods. Labels in black are shared species, and labels in red show the two non-shared, more abundant species per period.

Overall, Sørensen dissimilarity between past and present communities was high, indicating substantial species turnover. Specifically, total Sørensen dissimilarity was 0.70, of which 96% was attributable to species turnover (species replacement) and only 4% to nestedness (species loss/gain).

Lastly, the relative abundance of shared species also changed over time, reflecting shifts not only in the composition of shared species but also in which of them were more abundant in each period. For instance, the three most abundant species during the past period (1983-1984) were *Apis mellifera, Lasioglossum prasium,* and *Lasioglossum immunitum.* In the case of the present period (2020-2021), *Apis mellifera* was also the most abundant, followed by *Bombus terrestris, Xylocopa cantabrita,* and *Megachile sicula*.

## Discusion

The Doñana Natural Area is a cornerstone for pollinator conservation in the western Mediterranean, offering an exceptional range of habitat types shaped by dynamic climatic, hydrological, and geomorphological processes (Herrera, 1988). This structural and environmental heterogeneity sustains a high degree of functional and taxonomic diversity in its bee assemblages, making it a key refuge for wild pollinators at both regional and continental scales (Nieto et al., 2014). Although of great importance, bee diversity has never been systematically reported before in Doñana, along with additional information such as plant importance or historical trends.

### Composition of Bee Fauna in the Doñana Natural Area

We have created the first bee checklist with 385 bee species, 47 genera, and more than 27.802 plant-pollinator interaction observations. These findings reinforce the potential of biological surveys as an indispensable tool for understanding pollinator community dynamics and informing conservation strategies. The sustained effort to document bee diversity in Doñana—including recent initiatives to sample previously overlooked habitats—has substantially enriched our knowledge of local faunal composition.

Despite Doñana’s conservation significance, its pollinator communities are increasingly vulnerable to anthropogenic stressors. Hydrological alterations, land-use intensification, and climate-driven changes in resource availability have all contributed to shifts in habitat quality and community composition (Rodríguez-Rodríguez et al., 2020; AEMET & CSIC, 2022). Particularly concerning is the ongoing expansion of intensive agriculture around the area, accelerating aquifer depletion, reducing natural buffers, and increasing agrochemical exposure—factors that degrade essential nesting and foraging resources necessary for bees’ persistence (Arroyo et al., 2021; Green et al., 2024; Burkle, Marlin & Knight, 2013).

### Bee floral usage

We have established floral usage for 368 bee species out of the 382 total, resulting in a classification of plant importance that links plant importance in terms of floral resources with bee dependencies on those resources (Albrecht et al., 2012). These floral usages are aggregated observations across space and time, and bees could display distinct floral preferences over the season as floral communities change, or in response to competition and environmental stress. Among these stressors, competition with managed honey bees may play a significant role by altering the availability of floral resources and influencing wild bee foraging patterns (Magrach et al., 2017). For instance, some species consistently favor specific color cues or flower shapes, while others may adapt their preferences when preferred flowers are rare (Reverté et al., 2016). However, specialist bees are particularly vulnerable to changes as they lack the behaviour flexibility required to rewire their interactions, and they could experience steep population declines or local extinctions if their host plants decline due to land-use change, introduction of invasive species, or climate change (Williams et al., 2010; Burkle et al., 2013; Jacquemin et al., 2020).

We obtained the importance of plant species by combining the number of species they can sustain and how important they are for each species. Key species include abundant and attractive plants such as *Cistus crispus* or *Salvia rosmarinus*, which provide floral resources for a great number of species, many visiting exclusively this plant species in our dataset. Assessing plant importance and floral dependency helps identify keystone plants critical for conservation of bee communities. Keystone species are often common species that might support a greater number of pollinators and specialist species, promoting network resilience (Cagua et al., 2019). Previous studies have shown that the loss of a single keystone plant can disrupt the entire pollinator community, reducing interaction diversity and causing secondary losses in bees (Sandacz et al., 2023). Targeting these keystone species during restoration can help managers create more stable systems, enhancing pollinator diversity and ecosystem functioning (Rafferty et al., 2024).

### Historical trends

Our findings reveal a marked reorganization of wild bee assemblages in the Doñana Biological Reserve over the past four decades. Despite comparable levels of species richness between the historical (1984–1985) and recent (2020–2021) surveys, the community composition has shifted substantially, with only 15 of 78 total species shared across periods. This represents a temporal turnover of over 96%, indicative of significant ecological change.

The shifts in community composition are likely driven by environmental changes affecting species-specific habitat preferences, phenologies, and interactions with floral resources (Turley et al., 2022). However, we can not disentangle natural fluctuations from changes driven by environmental change. In any case, such a marked reorganization of bees’ assemblages is consistent with broader patterns of pollinator changes observed in other temperate climates (Burkle et al., 2013, Potts et al., 2010; Biesmeijer et al., 2006). The observed turnover may reflect the replacement of specialist species by opportunistic or disturbance-tolerant taxa such as *Megachile sicula* or *Eucera rufa*, a process that has been widely documented as ecosystems respond to anthropogenic pressures and climate variability (Clavel et al., 2011; Burkle et al., 2013). In Doñana, increasing environmental pressures—particularly drought intensification, land-use changes, and water mismanagement (Rodríguez-Rodríguez et al., 2021; AEMET & CSIC, 2022)—may be driving these shifts. These factors are likely to favor generalist and disturbance-tolerant species, while filtering out more specialised taxa with narrower habitat or floral requirements (Clavel et al., 2011; Burkle et al., 2013).

The observed community turnover may also reflect phenological shifts, where climate-induced changes in seasonal activity could alter detection patterns or disrupt bee–plant synchrony (Turley et al., 2022; Montes-Perez et al., 2025). However, our sampling design, constrained to a single phenological window in each survey period, limits our ability to distinguish phenological change from species turnover. Repeated intra-annual sampling would be needed to resolve these dynamics. Additionally, although both datasets stem from standardized protocols and taxonomic consistency was ensured, potential differences in sampling effort or microhabitat coverage cannot be entirely ruled out. Nonetheless, the large-scale compositional change observed indicates the dynamic nature of bee communities.

Another caveat relates to the inherent spatio-temporal variability of Mediterranean ecosystems. This, combined with high temporal and spatial variability in pollinator populations, means that even well-designed sampling frameworks may fall short of capturing the full ecological picture (Chacoff et al., 2018; Turley et al., 2022). Rarefaction analyses and species accumulation curves suggest that the observed diversity is likely an underestimate, reinforcing the need for long-term, multi-seasonal, and spatially stratified monitoring. Additionally, we are aware of potential spatial sampling bias, for example, with an underrepresentation of structurally complex or seasonally transient habitats such as dunes or marshlands. These areas often harbor specialized taxa and unique interactions that are especially sensitive to disturbance and environmental filtering (Clavel, Julliard & Devictor, 2011). Their inclusion is essential not only for documenting species richness but for assessing ecosystem resilience and predicting the consequences of biotic homogenisation under global change scenarios (Carvalheiro et al., 2013).

Ultimately, the conservation of wild bees in Doñana hinges on both robust biological monitoring and effective policy implementation. Efforts must extend beyond the protected core to encompass the broader socio-ecological landscape, addressing diffuse pressures such as hydrological stress and unsustainable agricultural expansion. As a flagship Mediterranean reserve, Doñana offers an unparalleled opportunity to integrate ecological research with biodiversity management. Continued and enhanced sampling, supported by collaborative frameworks and open-access data, will be essential to ensure the long-term protection of this pollinator-rich ecosystem.

## Conclusion

In summary, this study delivers the first comprehensive bee checklist for the Doñana Protected Area alongside a detailed overview of wild bee diversity, revealing notable species richness, temporal turnover, and key plant–pollinator associations crucial for conservation. Our findings underscore the importance of sustaining habitat heterogeneity and floral resource diversity, including specific plant species vital for pollination, while addressing water management and land-use pressures to safeguard pollinator communities. The observed dynamic shifts in community composition over recent decades raise concerns about the long-term stability of plant–pollinator networks under growing environmental stress. Therefore, integrating explicit pollinator conservation within protected-area management and establishing sustained monitoring regimes incorporating functional trait and network analyses will be essential to uphold the ecological integrity and resilience of Mediterranean ecosystems such as Doñana.

## Acknowledgments

The authors thank Alejandro Núñez Carvajal, Carlos M. Herrera, Denis Michez, Félix Torres, Guillaume Ghisbain, Luis Oscar Aguado, Simone Flaminio and Thomas J. Wood for helping with the identifications or confirmation of specimens. In particular, we thank T. J. Wood for his generous assistance and valuable advice regarding the taxonomy of the genus *Andrena*.

FPM and NMP thank Ainhoa Magrach, Oscar Godoy and José Manuel Herrera for constructive comments on the manuscript, which greatly improved its clarity and quality. We are deeply grateful to the administration of the Doñana Natural Space for granting research permits and for their constant support during fieldwork. We especially thank the park staff and rangers for their invaluable assistance in accessing and sampling remote areas, and for their dedication to the conservation of this unique ecosystem. FPM also acknowledges the collaboration of the Doñana Biological Station (EBD-CSIC), with special thanks to Guyonne Janss and Sofia Conradi for their unconditional support with permits and access keys, which greatly facilitated our work.

## Author’s contribution

**Francisco P. Molina**: conceptualization; data curation; investigation; methodology; writing -original draft preparation; writing – review & editing. **Nerea Montes-Pérez**: conceptualization; data curation; formal analysis; investigation; visualization; writing – review & editing. **Luis Villagarcía:** writing – review & editing. **Ignasi Bartomeus:** conceptualization; funding acquisition; writing – review & editing.

## Notes

Conflict of interests; The authors do not have conflict of interests.

### Competing Interest Statement

The authors have declared no competing interest.

